# The human leukemic oncogene MLL-AF4 promotes hyperplastic growth of hematopoietic tissues in *Drosophila* larvae

**DOI:** 10.1101/2022.11.08.515565

**Authors:** Julie A. Johannessen, Miriam Formica, Nora Rojahn Bråthen, Amani Al Outa, Miriam Aarsund, Marc Therrien, Jorrit M. Enserink, Helene Knævelsrud

## Abstract

*MLL*-rearranged leukemias are among the leukemic subtypes with poorest survival, and treatment options have barely improved over the last decades. Furthermore, despite increasing molecular understanding of the mechanisms behind these hematopoietic malignancies, this knowledge has had poor translation into the clinic. Identification of novel treatment methods is hampered by the lack of relevant *in vivo* models that allow for rapid identification of actionable drug targets and small molecule inhibitors. Here, we report a *Drosophila melanogaster* model system to explore the pathways affected in *MLL*-rearranged leukemia. We show that expression of the human leukemic oncogene MLL-AF4 in the *Drosophila* hematopoietic system resulted in increased levels of circulating hemocytes and an enlargement of the larval hematopoietic organ, the lymph gland. Strikingly, depletion of *Drosophila* orthologs of known interactors of MLL-AF4, such as DOT1L, rescued the leukemic phenotype. In agreement, treatment with small-molecule inhibitors of DOT1L also prevented the MLL-AF4-induced leukemia-like phenotype. Taken together, this model provides an *in vivo* system to unravel the genetic interactors involved in leukemogenesis and offers a strategy for a prompt identification of potential therapeutic options for treatment of *MLL*-rearranged leukemia.

## Introduction

*MLL*-rearranged leukemias are associated with dismal overall outcome, with 5-year overall survival rates as low as 8-20% depending on the subtype (Issa et al., 2021; Richard-Carpentier et al., 2021). *MLL*-rearranged leukemia is characterized by cytological alterations of chromosome 11q23, which were first identified in a wide range of patients with both myeloid and lymphocytic leukemia, and later mapped to the *MLL* gene (Djabali et al., 1992; Gu et al., 1992; Tkachuk, Kohler, & Cleary, 1992; Ziemin-van der Poel et al., 1991). The *MLL* gene encodes a methyltransferase and has therefore been renamed *Lysine [K]-specific Methyltransferase 2A* (*KMT2A*). Despite the efforts to improve and develop new treatment over the last 50 years, little progress has been made and patients diagnosed with *MLL*-rearranged leukemia are left with a sparse range of therapy options and suffer poor survival rates.

Normal MLL positively regulates gene expression through histone modifications, directly as a histone 3 lysine 4 (H3K4) methyltransferase and indirectly by acting as an activator of the histone 4 lysine 16 (H3K16) acetyltransferase MOF (Dou et al., 2005). Among the main target genes of normal MLL are *homeobox* (*Hox)* genes which specify regions of the body plan through development and are involved in haematopoietic development (Yu, Hess, Horning, Brown, & Korsmeyer, 1995), and *Meis* genes (Zeisig et al., 2004). Regulation of both sets of genes are aberrant in leukemic patients carrying *MLL* chromosomal rearrangement (Armstrong et al., 2002; Rozovskaia et al., 2001). In general, the N-terminal part of MLL is responsible for identifying and binding the target gene sequences. This occurs mainly through interaction with Menin, which recruits the H3K36 di/trimethylation reader LEDGF (A. Yokoyama & Cleary, 2008; Akihiko Yokoyama et al., 2005; A. Yokoyama et al., 2004; Zhu et al., 2016), and a CxxC domain, which binds to unmethylated CpG sequences (Bina et al., 2013; Birke et al., 2002). The C-terminal part of MLL contains the MOF interaction site and the SET domain responsible for the histone H3 lysine 4 methylation, associated with normal regulation of the target genes (Milne et al., 2002).

In the MLL fusion proteins resulting from chromosomal translocations involving the *MLL* gene, the N-terminal part of MLL including the CxxC domain is retained, whereas the remainder of the protein is replaced by the C-terminal part of the fusion partner. To date, more than 100 such fusion partners have been described (Meyer et al., 2018), but the most common ones represent about 80% of clinical cases (MLL fused with either AF4/AFF1, ENL/MLLT1, AF9/MLLT3, AF10/MLLT10 or ELL) and exist in related complexes involved in regulation of transcriptional elongation (Slany, 2020). For instance, the fusion partner *ALL1-fused gene from chromosome 4* (AF4) is normally a nuclear protein involved in transcriptional activation (Li, Frestedt, & Kersey, 1998), which directly interacts with the histone acetylase ENL (Mueller et al., 2007). The MLL-AF4 fusion protein is thought to drive a feed-forward loop resulting in high activity of the histone H3 lysine 79 methyltransferase DOT1L, culminating in aberrant transcription of target genes (Slany, 2020). Even in the presence of this molecular understanding of the impact of MLL-fusion proteins, therapies developed towards these targets have yet to successfully enter clinical practice. In this context, additional genetically tractable animal models that allow rapid and unbiased *in vivo* genetic analysis and drug screens could complement studies in existing mouse models.

*Drosophila* has a long tradition as a system to understand gene function and to model disease, including cancer and leukemia (Al Outa, Abubaker, Madi, Nasr, & Shirinian, 2020). *Drosophila* hematopoiesis has several key similarities to mammalian hematopoiesis, as it is governed by conserved transcriptional programs that result in generation of hemocytes with functional similarity to human blood cells (Boulet, Miller, Vandel, & Waltzer, 2018). Fly immune cells reside in specific compartments where they interact with the microenvironment (Banerjee, Girard, Goins, & Spratford, 2019). Briefly, hematopoiesis in *Drosophila* occurs in two waves, first in the embryo and later in the larval hematopoietic organ, the lymph gland (Holz, Bossinger, Strasser, Janning, & Klapper, 2003; Jung, Evans, Uemura, & Banerjee, 2005). The lymph gland consists of two primary (anterior) and several secondary (posterior) lobes. The primary lobe can be divided into the progenitor-containing medullary zone, the cortical zone containing differentiated hemocytes and the posterior signaling center (PSC), which is involved in regulating the balance between progenitors and differentiated hemocytes (Kharrat, Csordás, & Honti, 2022). Traditionally, hematopoietic differentiation from prohemocytes has been known to result in three main types of hemocytes: Plasmatocytes, crystal cells and lamellocytes. The macrophage-like plasmatocytes make up the majority of hemocytes (around 95% of the hemocyte population) with a smaller fraction of crystal cells (around 5%) involved in melanization and wound healing (Honti, Csordás, Kurucz, Márkus, & Andó, 2014). Finally, large, actin-rich lamellocytes develop upon specific insults, such as the presence of parasitic wasp eggs, and are not normally observed in healthy larvae (Lanot, Zachary, Holder, & Meister, 2001; Rizki & Rizki, 1992). Recently, novel single cell sequencing data has unraveled a more diverse range of hematopoietic cell subtypes, including intermediate stages (Cho et al., 2020; Girard et al., 2021). Furthermore, hemocyte development is not limited to the linear development from prohemocytes as a common precursor. There are also mechanisms of transdifferentiation, where plasmatocytes can develop into lamellocytes, underpinning the plasticity and intricacy of the *Drosophila* hematopoietic system (Honti et al., 2010).

Notably, the MLL protein was first identified as Trithorax (Trx) in *Drosophila melanogaster* (Schuettengruber, Bourbon, Di Croce, & Cavalli, 2017), and orthologs of all the main MLL fusion partners are found in *Drosophila*. Human full-length MLL was previously found to partly rescue a *trx* mutant. Interestingly, expression of human MLL-AF4 or MLL-AF9 in *Drosophila* larval brain affected cell cycle progression, whereas ubiquitous expression was lethal (Muyrers-Chen et al., 2004). Furthermore, expression of human MLL-AF10 was found to deregulate the activity of *Polycomb* group-responsive elements in a *Drosophila* adult eye reporter system (Perrin et al., 2003), indicating that these human leukemic fusion proteins are active also in *Drosophila*.

In this study, we exploited the conservation of the MLL complex and hematopoiesis in *D. melanogaster* to gain insight into the hematopoietic malignancies that occur upon chromosomal *MLL*-rearrangements. We show that expression of human MLL-AF4 in the *Drosophila* hematopoietic system leads to increased levels of differentiated hemocytes, resulting in enlarged lymph glands and enhanced numbers of circulating hemocytes. These phenotypes were dependent on the main complex partners that have also been identified in human leukemia and could be rescued by genetic or chemical targeting of DOT1L.

## Results

### Expression of the human oncogene MLL-AF4 induces a leukemia-like phenotype in flies

Due to the conserved function of the equivalent of the human MLL complex in *Drosophila melanogaster*, the trithorax complex, we hypothesized that expressing the human oncogene MLL-AF4 in the hematopoietic system of *D. melanogaster* could induce a leukemic phenotype. We expressed either human full-length MLL or the human fusion protein MLL-AF4 in the larval hematopoietic system using the *pxn*-Gal4 driver and co-expressing GFP (*pxn*- Gal4>UAS-GFP). The *pxn*-Gal4 driver was chosen because *peroxidasin* is expressed in the cortical zone of the lymph gland from the second instar larval stage and onwards in the maturing hemocytes of the plasmatocyte and crystal cell lineages (Jung et al., 2005; Nelson et al., 1994). We assessed the overall effects of MLL-AF4 expression on the lymph gland by measuring lymph gland size changes. Furthermore, we evaluated the effects of MLL-AF4 expression on the cortical zone by quantification of the GFP+ volume fraction of the lymph gland, to identify whether this compartment changed in size relative to the overall lymph gland. Interestingly, while the lymph glands in larvae expressing full-length, normal MLL in the cortical zone were similar to wild-type (WT) lymph glands, expression of the MLL-AF4 oncogene induced significant lymph gland enlargement and an increase in GFP+ hemocytes in the lymph gland compared to WT larvae (Fig 1A-G). Furthermore, lymph glands expressing MLL-AF4 in the cortical zone displayed increased levels of GFP volume per total lymph gland volume (Fig 1H). Flow cytometry of dissociated lymph glands also showed an increase in the total number of GFP+ hemocytes per lymph gland as well an increase in the percentage of GFP+ hemocytes per lymph gland (Fig 1I-J). Lymph glands expressing either the MLL N-terminal or the AF4 C-terminal part of the MLL-AF4 fusion protein in the cortical zone were similar to controls (Supp. Fig 1A-F). We verified by RT-qPCR that the MLL N-terminal or the AF4 C-terminal portions were expressed to similar levels as the MLL-AF4 fusion protein (Supp. Fig 1G-I). We also found that simultaneous expression of two independent UAS-driven MLL-AF4 transgenes induced even larger lymph glands relative to a single MLL-AF4 transgene, bolstering the causal effect of MLL-AF4 expression on the hyperplasia phenotype (Supp. Fig 1J-L). To investigate whether the increase in lymph gland size was a result of increased proliferation, we stained lymph glands from L2, feeding L3 and wandering L3 stages for the proliferative marker phosphorylated histone 3 serine 10 (pH3Ser10). Indeed we found that the lymph gland overgrowth could be explained by increased proliferation in MLL-AF4-expressing lymph glands at the L2 larval stage, whereas there was no difference in proliferation in lymph glands from feeding and wandering L3 larvae (Supp. Fig 2A-G).

**Figure 1:**
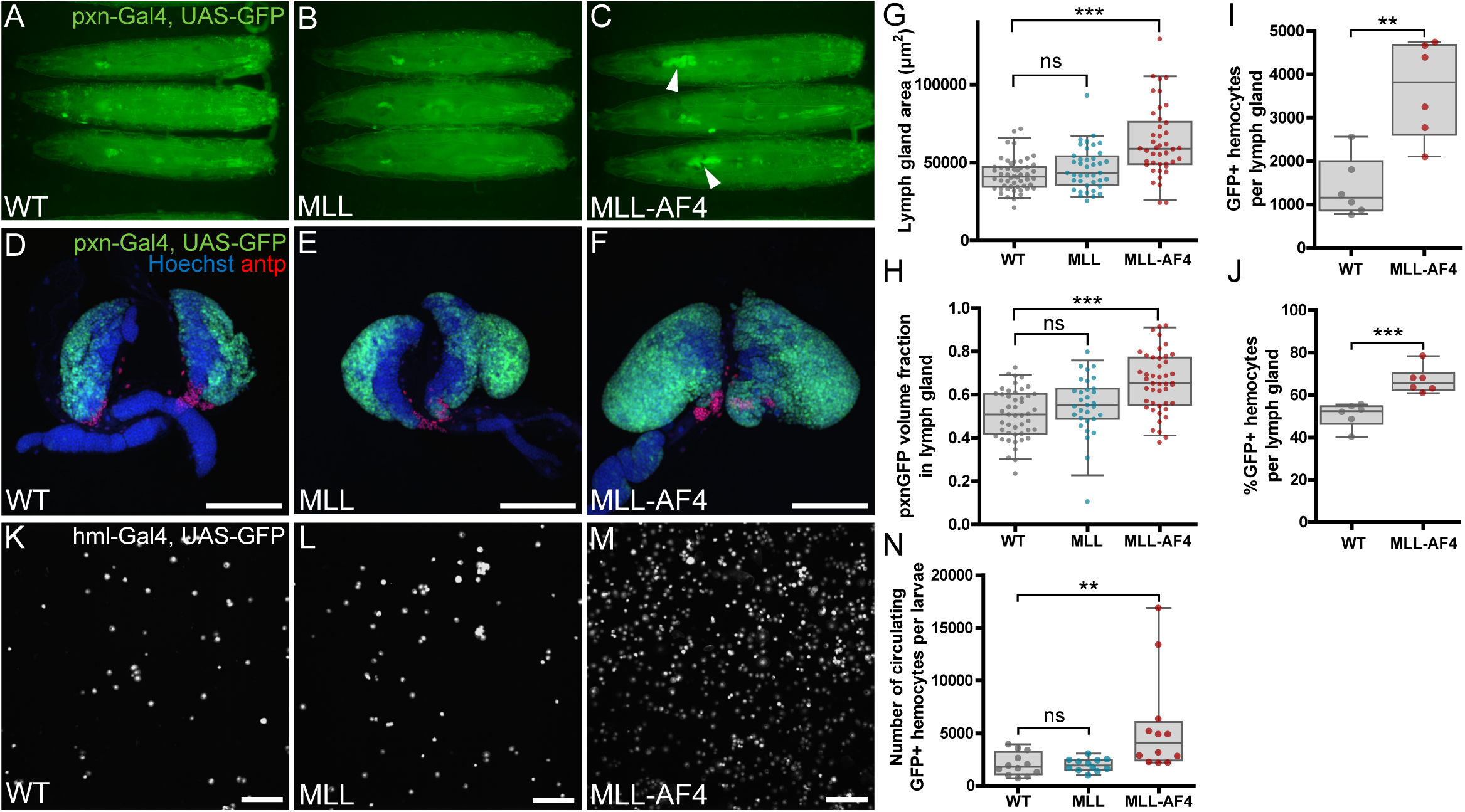
Expression of the human oncogene MLL-AF4 in the hematopoietic system of *D. melanogaster* induces a leukemia-like phenotype. **A-C:** Representative images of wandering third instar larvae expressing full-length human MLL or human MLL-AF4 driven by *pxn*-Gal4 and imaged by widefield fluorescence microscopy. The hematopoietic system is marked by GFP expression driven by *pxn*-Gal4. Arrowheads indicate enlarged lymph glands. **D-F:** Immunofluorescence confocal images of lymph glands that are WT or expressing MLL or MLL-AF4 driven by *pxn*-Gal4. The cortical zone is marked by GFP, and the posterial signaling center (PSC) is detected by immunostaining against antp (in red). Images are maximum intensity projection of Z-stacks. Scale bars 100 µm. **G:** Quantification of lymph gland area (µm^2^) from immunofluorescence confocal images. **H**: Quantification of GFP-positive lymph gland volume presented as ratio of total lymph gland volume based on immunofluorescence confocal stacks. **I**: Quantification of GFP positive hemocytes per lymph gland by flow cytometry. **J**: Quantification of percentage of GFP-positive cells in larval lymph glands measured by flow cytometry. Graph shows combined results from around 15 lymph glands per genotype from 6 independent experiments. **K-M:** Immunofluorescence confocal images of circulating hemocytes from larvae that are WT or expressing MLL or MLL-AF4 driven by hml-Gal4. Images are maximum intensity projection of Z-stacks. Scale bars 100 µm. **N:** Quantification of number of circulating hemocytes expressing GFP. Values are per larva across 4 larvae in 3 replicates. **G, H, N:** Error bars show the 95^th^ percentile. Statistical significance was determined by one-way ANOVA with Bonferroni post-test to assess significant differences from WT. **I, J:** Error bars show the 95^th^ percentile. Statistical significance was determined by two-sided Student’s t-test to assess significant differences from WT. **Genotypes: A, D, G-J**: *pxn-Gal4, UAS-GFP/+* (WT) **B, E, G-J:** *pxn-Gal4, UAS-GFP/UAS-MLL* **C, F, G-J:** *pxn-Gal4, UAS-GFP/UAS-MLL-AF4* **K, N**: *hml-Gal4, UAS-GFP/+* (WT) **L, N:** *hml-Gal4, UAS-GFP/UAS-MLL* **M, N:** *hml-Gal4, UAS-GFP/UAS-MLL-AF4*

**Figure 2:**
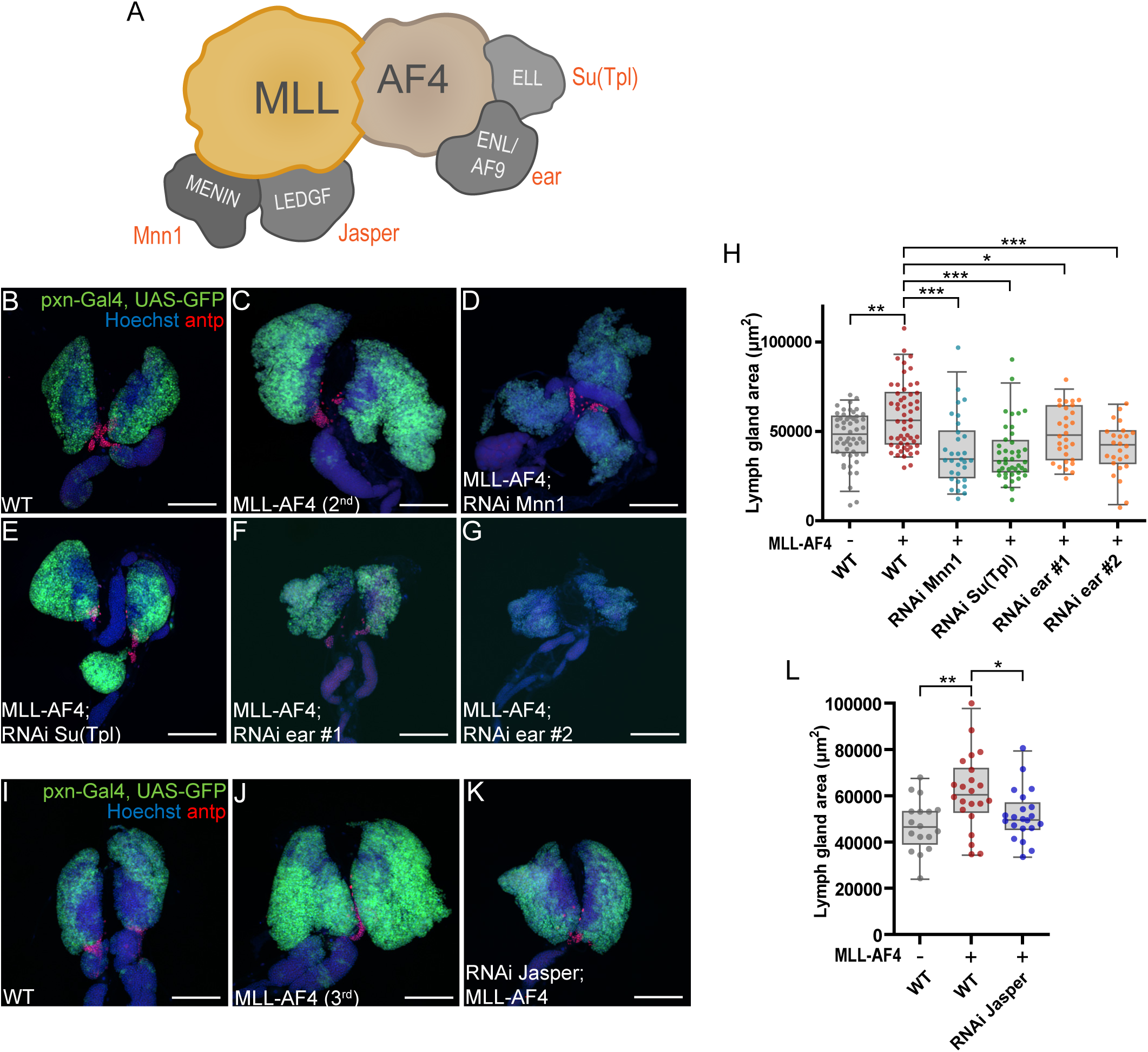
MLL-AF4-driven lymph gland enlargement depends on *Drosophila* homologs of known MLL-AF4 interaction partners. **A**: Schematic overview over some of the known direct interactors with the fusion protein MLL-AF4 in humans based on current literature. The corresponding *D. melanogaster* homologues are indicated in red. **B-G and I-K**: Representative immunofluorescence confocal images of lymph glands with *pxn*-Gal4-driven expression of MLL-AF4 and concurrent depletion of indicated target genes (**B-G**: MLL-AF4 on the 2^nd^ chromosome, **I-K**: MLL-AF4 on the 3^rd^ chromosome). The cortical zone is marked by GFP expression and PSC is detected by immunostaining for antp (in red). Images are maximum intensity projection of Z-stacks. Scale bar 100 µm. **H and L:** Quantification of lymph gland area from immunofluorescence images with MLL-AF4 expression driven by pxn-Gal4 combined with RNAi (**H**: MLL-AF4 on the 2^nd^ chromosome, **M**: MLL-AF4 on the 3^rd^ chromosome). **H, L**: Error bars show 95^th^ percentile. One-way ANOVA with Bonferroni post-test was performed to show significant differences from MLL-AF4. **Genotypes: B, H, I, L:** *pxn-Gal4, UAS-GFP/+* (WT) **C, H:** *pxn-Gal4, UAS-GFP/UAS-MLL-AF4* **D, H:** *pxn-Gal4, UAS-GFP/UAS-MLL-AF4; UAS-RNAi Mnn1 GL00018/+* **E, H:** *pxn-Gal4, UAS-GFP/UAS-MLL-AF4; UAS-RNAi Su(Tpl) HMS00277/+* **F, H:** *pxn-Gal4, UAS-GFP/UAS-MLL-AF4; UAS-RNAi ear HMS00107/+* **G, H:** *pxn-Gal4, UAS-GFP/UAS-MLL-AF4; UAS-RNAi ear JF02905/+* **J, L:** *pxn-Gal4, UAS-GFP/+; UAS-MLL-AF4/+* **K, L:** *pxn-Gal4, UAS-GFP/UAS-RNAi Jasper HMC03961; UAS-MLL-AF4/+*

Given that *pxn-*Gal4 mainly drives MLL-AF4 expression in differentiating hemocytes (Jung et al., 2005), we determined whether its expression affected hemocyte maturation. The observed increase of GFP+ hemocytes was identified as plasmatocytes as shown by staining the primary lobes for NimC1/P1(Kurucz, Markus, et al., 2007) (Supp. Fig 3A-D). Furthermore, crystal cell levels were increased upon MLL-AF4 expression as evaluated by staining for Hindsight (Hnt) (Terriente-Felix et al., 2013) (Supp. Fig 3E-H). Finally, lamellocytes were neither observed in control glands nor in MLL-AF4 expressing lymph glands (Supp. Fig 3I-K). Thus, expression of MLL-AF4 induces aberrations in the hematopoietic development of *D. melanogaster*, leading to increased numbers of differentiated hemocytes.

**Figure 3:**
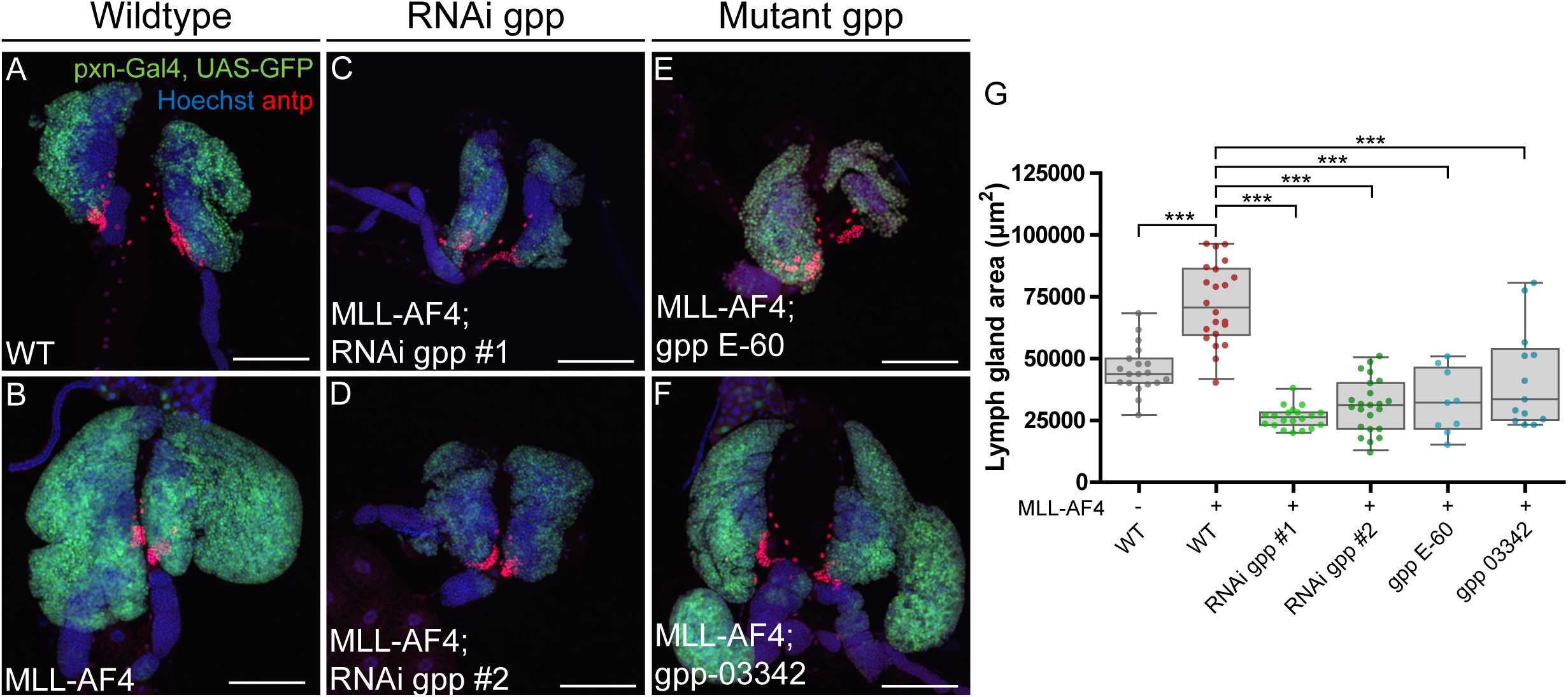
MLL-AF4-driven lymph gland enlargement depends on the *Drosophila* homolog of DOT1L, gpp. **A-F**: Representative immunofluorescence confocal images of lymph glands depleted of *gpp* in combination with MLL-AF4 expression driven by pxn-Gal4. The cortical zone is marked by GFP expression and PSC is detected by immunostaining for antp (in red). Images are maximum intensity projection of Z-stacks. Scale bars 100 µm. **A, B**: WT lymph gland and UAS-MLL-AF4 expressing lymph gland. **C, D**: MLL-AF4 expressing lymph glands combined with expression of dsRNA against *gpp*. **E, F**: MLL-AF4 expressing lymph glands from a heterozygous genetic background containing *gpp* mutants E-60 or 03342. **G:** Quantification of lymph gland area (µm^2^) from immunofluorescence confocal images. Error bars show standard deviation. One-way ANOVA with Bonferroni post-test was performed to show significant differences from MLL-AF4. **Genotypes: A, G:** *pxn-Gal4, UAS-GFP/+* (WT) **B, G:** *pxn-Gal4, UAS-GFP/UAS-MLL-AF4* **C, G:** *pxn-Gal4, UAS-GFP/UAS-MLL-AF; UAS-RNAi gpp HMS00160/+* **D, G:** *pxn-Gal4, UAS-GFP/UAS-MLL-AF4; UAS-RNAi gpp JF1283/+* **E, G:** *pxn-Gal4, UAS-GFP/UAS-MLL-AF4; UAS-gpp E-60/+* **F, G:** *pxn-Gal4, UAS-GFP/UAS-MLL-AF4; UAS-gpp 03342/+*

To further understand how MLL-AF4 affects the larval hematopoietic system, we investigated the effect of expressing MLL-AF4 in circulating hemocytes using the *hml*-Gal4 driver and co-expressing GFP (*hml*-Gal4>UAS-GFP) (Goto, Kadowaki, & Kitagawa, 2003). Whereas larvae expressing full-length MLL had normal levels of circulating hemocytes, MLL-AF4 expression resulted in a significant increase in circulating hemocyte levels (Fig 1K-N). We also noticed that in larvae expressing MLL-AF4, lamellocytes were specifically present in the hemolymph, which is indicative of aberrant hemocyte differentiation (Supp. Fig 4C and D). Taken together, these results show that MLL-AF4 induces enlargement of hematopoietic tissues in *Drosophila* primarily through an increase in plasmatocytes. In addition, MLL-AF4 expression affects hemocyte fate since it promotes the development of lamellocytes.

**Figure 4:**
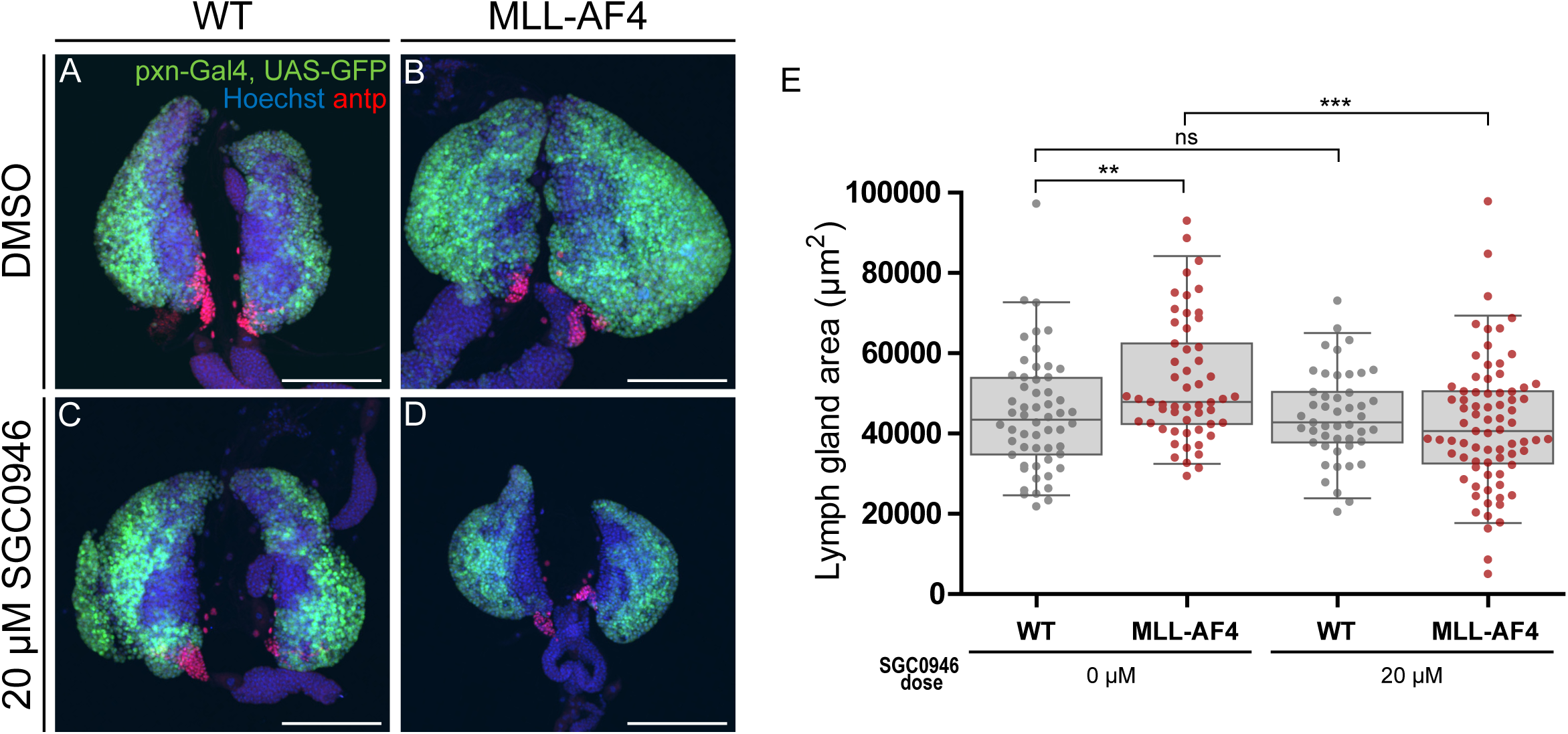
The MLL-AF4 *Drosophila* model is sensitive to DOT1L drug treatment. **A-D:** Lymph glands from wandering 3^rd^ instar larvae. Cortical zone is marked by GFP expression and PSC is detected by immunostaining for antp (in red). Images are maximum intensity projection of Z-stacks. Scale bars 100 µm. **A, B:** WT and MLL-AF4 expressing lymph glands treated with DMSO only. **C, D:** WT and MLL-AF4 expressing lymph glands treated with 20 µM SGC0946. **E:** Quantification of lymph gland area (µm^2^) from immunofluorescence confocal images from drug treated larvae with either DMSO or the DOT1L inhibitor SGC0946. Error bars of box plots show the 95^th^ percentile. Statistical significance was determined by two-sided Student’s t-test to assess significant differences between drug treated larvae and DMSO control for each genotype. **Genotypes: A, C, E**: *pxn-Gal4, UAS-GFP/+* (WT) **B, D, E:** *pxn-Gal4, UAS-GFP/UAS-MLL-AF4*

### Knockdown of MLL-AF4 complex partners rescues the leukemic phenotype

Next, we tested whether the observed leukemia-like phenotypes depend on fly orthologs of the known complex partners of the human MLL-AF4 fusion protein (see Figure 2A for a schematic overview). To this end, we used RNAi-mediated knockdown of these orthologs. For MLL interaction partners we depleted Mnn1 (the *Drosophila* ortholog of MEN1) or Jasper (the *Drsophila* ortholog of LEDGF/PSIP1), since the human orthologs are required for MLL to interact with chromatin (Fig 2A). For interaction partners of human AF4, we targeted Su(Tpl) (the *Drosophila* ortholog of ELL) and ear (the *Drosophila* ortholog of ENL/MLLT1 and AF9/MLLT3), which are part of the transcriptional super elongation complex (SEC) (Luo, Lin, & Shilatifard, 2012) (Fig 2A). Knockdown efficiency was evaluated by RT-qPCR in hemocytes for several available RNAi lines against each of these targets (Supp Table 1 and Supp. Fig 5A-D) and a subset of lines producing efficient knockdown was selected for phenotypic analysis. *Pxn*-driven depletion of most of these gene products in the cortical zone of otherwise wild-type lymph glands did not result in obvious changes in lymph gland size or morphology (Supp. Fig 5E-L). The exception was depletion of *Mnn1*, where lymph glands presented with larger secondary lobes than the WT, and the overall lymph gland size was smaller than WT (Supp. Fig 5G, L). Interestingly, in lymph glands expressing MLL-AF4 in the cortical zone, concurrent depletion of *Mnn1*, *Jasper*, *ear* or *Su(Tpl)* resulted in a reduction in overall lymph gland size (Fig 2B-L). Taken together, this shows that the observed leukemia-like phenotype of MLL-AF4 depend on the same conserved interaction partners as in human leukemia.

### The leukemic phenotype of the *Drosophila* MLL-AF4 model depends on the DOT1L fly homolog *gpp*

In *MLL*-rearranged leukemia, the histone methyltransferase DOT1L has been shown to be associated with the MLL fusion protein complex and DOT1L is required for leukemogenesis in different cellular and organismal model systems (Bernt et al., 2011; Mohan et al., 2010). Consequently, several clinical studies have focused on DOT1L as a potential therapeutic target, mainly in combination therapy with standard treatment (Klaus et al., 2014; Menghrajani et al., 2019; Stein et al., 2018). To test whether DOT1L is required for the MLL-AF4 phenotype in *Drosophila*, we first depleted the *Drosophila* DOT1L homolog *gpp* (Shanower et al., 2005) in lymph glands expressing MLL-AF4 in the cortical zone. Interestingly, *gpp* knockdown using two different RNAi lines resulted in complete rescue of the MLL-AF4-driven increase in lymph gland size (Fig 3A-D, G). A similar rescue of the leukemic phenotype was also observed when MLL-AF4 was expressed in a heterozygous genetic background with a mutant allele of *gpp* (*gpp E-60* or *gpp-03342*) (Fig 3E-G), in which *gpp* expression is reduced by half (Supp. Fig 6H). In contrast, in WT larvae, knockdown of *gpp* or introduction of *gpp* mutant alleles did not result in a significant reduction of lymph gland size (Supp Fig 6A-F). Reduction of *gpp* expression in hemocytes by RNAi or mutant alleles was verified by RT-qPCR (Supp. Fig 6G-H). Altogether, these data show that *Drosophila* DOT1L is essential for the phenotype induced by human MLL-AF4 in this *in vivo* model system.

### The MLL-AF4 leukemia phenotype can be rescued by DOT1L inhibitors

To understand if this model offers a reliable and rapid platform for identifying pharmacological inhibitors for human MLL-r leukemia, we next addressed whether drugs in pre-clinical and early-stage clinical studies are efficacious in the *Drosophila* MLL-AF4 leukemia model. We selected inhibitors targeting different downstream mechanisms of MLL-AF4. Specifically, we tested the inhibitor of MLL-menin interaction MI-463, the BRD inhibitor i-BET and the DOT1L inhibitor SGC0946. Drugs of interest were administered to developing larvae by mixing them into the standard fly food. Drug effect on the hematopoietic system was assessed in wandering 3^rd^ instar larvae. Interestingly, initial experiments assessing drug effects by visualizing the hematopoietic system through the cuticle of intact larvae revealed that treatment with the DOT1L inhibitor seemingly reduced lymph gland size (Supp. Fig. 7A-L), whereas MI-463 had little effect and i-BET induced larval death (Supp. Fig 7M-T). Therefore, we decided to investigate the DOT1L inhibitor effect further. Larvae were treated as before, and lymph glands were dissected out to assess drug effects on lymph gland size and morphology. Treatment with 20 µM SGC0946 was shown to fully rescue the lymph gland hyperplasia phenotype observed in the leukemia model (Fig 4A-F). Notably, a subset of SGC0946-treated MLL-AF4-expressing lymph glands were even smaller than WT lymph glands, indicating that MLL-AF4-expressing hemocytes are particularly sensitive to inhibition of gpp/DOT1L. Together, these data provide proof-of-principle that the MLL-AF4 *Drosophila* model can be used for *in vivo* drug screens to identify compounds that inhibit leukemogenesis and to explore the epigenetic landscape of leukemogenesis.

## Discussion

*MLL*-rearranged leukemia is associated with poor overall survival, and there exists a need for *in vivo* models to identify actionable targets and novel compounds for treatment of this type of leukemia. Here, we characterized a novel *Drosophila* model, where expression of the human leukemic oncogene MLL-AF4 results in a leukemia-like phenotype that can be rescued by genetic or pharmacologic targeting of components required for leukemia development in mammalian model systems of *MLL*-rearranged leukemia. This low-cost and rapid-to-screen model with the associated comprehensive toolbox for genetic manipulation and unbiased screening will allow for novel discoveries and sought-after therapeutic options *in vivo*.

In mouse models, DOT1L has been studied as a potential therapeutic target in *MLL*-rearranged leukemia, using inhibitors that either inhibit its catalytic activity or that disrupt its interaction with MLL fusion partners (Wu et al., 2021). Multiple mechanisms have been proposed to explain how the methyltransferase DOT1L contributes to leukemogenesis. Traditionally, it is believed that the fusion partner homologues AF9 and ENL are responsible for recruitment of DOT1L through the multi-subunit complex EAP (ENL-associated proteins), but DOT1L has also been shown to be directly associated with AF10 (Slany, 2016). Furthermore, although MLL-AF4 does not appear to be directly associated with DOT1L, it can stimulate DOT1L activity indirectly through the recruitment of AF9 (Biswas et al., 2011; Lin et al., 2010). Together with our findings that DOT1L inhibitors rescue the MLL-AF4 phenotype, these studies underscore DOT1L as a relevant drug target for different MLL fusion proteins. Pharmacological inhibition of DOT1L with small molecules selectively kills *MLL*-rearranged leukemia cells *in vitro* and *in vivo* (Scott R. Daigle et al., 2013; S. R. Daigle et al., 2011). In line with this, we find that a subset of MLL-AF4-expressing lymph glands from larvae raised on SGC0946-containing food are even smaller than WT lymph glands, indicating that MLL-AF4-expressing hemocytes are particularly sensitive to pharmacological gpp/DOT1L inhibition. Therefore, this fly model of leukemia could also be used as a simplified *in vivo* screening tool for developing DOT1L inhibitors or identifying associated pathways.

We also found genetic depletion of *gpp/DOT1L* to rescue the leukemia-like phenotype of our *MLL*-rearranged leukemia model. Interestingly, the *gpp* allele E-60 was identified as an enhancer of the phenotype caused by expression of the human leukemic oncogene NUP98-HOXA9 in the *Drosophila* eye (Gavory et al., 2021). In contrast, we find this allele to suppress the phenotype in our larval model of *MLL*-rearranged leukemia, similar to the *gpp* allele 03342, which previously failed to complement lethal *gpp* alleles (Shanower et al., 2005). Cells transformed by mechanistically distinct leukemic oncogenes, such as MLL-AF9, MLL-AF6 and NUP98-NSD1, have all shown sensitivity to DOT1L inhibition (Deshpande et al., 2013; Deshpande et al., 2014; Nguyen, Taranova, He, & Zhang, 2011). As these oncogenes are known to activate *HOX* genes, DOT1L is probably promoting aberrant transcription of the *HOX* gene cluster in these malignancies. When it comes to NUP98-HOXA9, the lack of gpp/DOT1L dependency might be due to the fusion oncoprotein containing the downstream target itself, hence bypassing the regulative effect of DOT1L on *HOX* genes. In this case, based on the *Drosophila* model (Gavory et al., 2021), gpp/DOT1L appears to rather oppose NUP98-HOXA9 function, underscoring the need to critically evaluate the use of DOT1L inhibitors across cytogenetically different subtypes of leukemia.

Menin and LEDGF are other potential targets in MLL-r leukemia where small molecule inhibitors have been developed (Borkin et al., 2015; Grembecka et al., 2012; He et al., 2016), and genetic depletion of the *Drosophila* orthologs *Mnn1* and *Jasper* suppressed the lymph gland enlargement in our MLL-AF4 fly model. In this model, the small molecule MI-463, which inhibits MEN1-MLL interaction, only produced a modest effect on the MLL-AF4-induced lymph gland hyperplasia. This could be due to limited bioavailability of the compound, or to differences in how *Drosophila* Mnn1 interacts with MLL-AF4 compared to human MEN1. Future studies, that test larger concentration ranges of other inhibitors of MEN1 activity or other MEN1-MLL interaction inhibitors, could address this concern. Furthermore, inhibitors of the bromodomain and extra-terminal (BET) family of proteins such as i-BET have shown promising results in preclinical models and in early clinical trials on hematologic malignancies (Abedin, Boddy, & Munshi, 2016), although clinical implementation of most of these inhibitors has been hampered by toxicity (Shorstova, Foulkes, & Witcher, 2021). Indeed, larvae feeding on food containing i-BET died at early larval stages irrespective of the presence of MLL-AF4, indicating general toxicity at concentrations above 5 µM. Therefore, we believe that our *Drosophila* model can be used to quickly discard candidate compounds or derivatives that are too toxic for further clinical development, which may help to reduce attrition commonly associated with drug development (Waring et al., 2015).

Although previous *MLL*-rearranged leukemia model systems have been reported in *D. melanogaster*, they have been limited to the observation of lethality, cell cycle defects (Muyrers-Chen et al., 2004) or eye phenotypes in a reporter-assay of *Polycomb* group-responsive element activity (Perrin et al., 2003). These studies showed that expression of human MLL-AF4, MLL-AF9 or MLL-AF10 affected cell cycle progression, normal development or transcriptional regulation. In contrast, our viable *in vivo* model allows for a more detailed investigation of the hematopoietic irregularities that are induced upon MLL-AF4 expression. We observed changes in levels of differentiated hemocytes, specifically increased levels of plasmatocytes and crystal cells in the lymph gland as well as the presence of lamellocytes in circulation. These results are qualitatively more similar to the larval leukemia model based on NUP98-HOXA9 expression (Baril et al., 2017).

In broader terms, *Drosophila* has previously been successfully employed to model leukemia-like phenotypes of other human leukemic oncogenes, such as AML1-ETO, NUP98-HOXA9 and BCR-ABL (Abubaker et al., 2022; Baril et al., 2017; Breig et al., 2014; Fogerty et al., 1999; Osman et al., 2009; Outa et al., 2020; Sinenko et al., 2010). In the context of BCR-ABL, a fly model of BCR-ABL expression in the fly eye has been put forward as a screening platform for identification of novel or improved compounds (Outa et al., 2020). In our study we find that the larval aberrant hematopoietic phenotype induced by human MLL-AF4 expression can be suppressed by DOT1L inhibitors, indicating that our model could be used to test derivatives of current inhibitors or novel compounds. The phenotype is amenable to high-throughput imaged-based screening, similar to what has previously been performed for a *Drosophila* Ras-driven tumor model (Willoughby et al., 2013), as the enlarged GFP-positive lymph gland can be visualized through the semi-transparent larval cuticle.

Collectively, we present a genetically tractable MLL-rearranged leukemia model that is sensitive to drug screening and that can be exploited to further investigate the role of MLL-AF4 in leukemogenesis.

## Materials and methods

### Fly stocks

Fly stocks were mainly obtained from Bloomington Stock Center or VDRC. *UAS-MLL-AF4* ML5 (on 2nd), *UAS-MLL-AF4* ML6 (on 3rd), *UAS-MLL* FL on 2^nd^, *UAS-MLL* FL on 3^rd^, *UAS-MLL N-term* (on 3rd), *UAS-MLL N-term* (on X) and *UAS-AF4 C-term* (on X) were reported elsewhere (Muyrers-Chen et al., 2004). A full list of the fly lines used in this study can be found in Supplementary Table 1. All crosses were performed on standard potato mash fly food (27.3 g/liter dry yeast, 32.7 g/liter dried potato powder, 60 g/liter sucrose, 0.73% agar, 0.2% 4-hydroxybenzoic acid methyl ester (nipagin)/ethanol and 0.45% propionic acid) with additional yeast paste and incubated at 25 °C for the first 72 hrs followed by 48-72 hrs at 29 °C to boost Gal4-driven expression.

### GFP imaging of entire larvae

Larvae were heat fixed to visualize the lymph gland in intact animals. Wandering L3 larvae were collected, washed once in PBS with 0.2% Triton-X100 (Sigma Aldrich, Cat#T9485) (PBT 0.2%) and once in PBS, followed by heat fixation in a drop of glycerol on a microscope slide on a heat block set at 70 °C for approximately 10 seconds. A coverslip was added on top of the glycerol drop with the heat fixed larvae before imaging them with a GFP filter on Leica MZFLIII stereomicroscope and LAS v4.9 software. Post-imaging enhancement of GFP signal for sufficient visualization was performed in Adobe Photoshop.

### Immunofluorescence confocal microscopy of lymph glands

For dissection of lymph glands, larvae were washed once in PBT 0.2% and once in PBS 1x. Lymph glands were dissected in PBS 1x together with the mouth hook and CNS to simplify washing and staining. The tissue was fixed with 4% paraformaldehyde (Polysciences, Cat# 18814-20) for 10 min, and then washed in PBT 0.2% three times. Samples were then blocked in PBT 0.2% with 5% bovine serum albumin (Roche, Cat# 10735094001) before staining with primary antibodies overnight at 4 °C. The primary antibodies used in this study were Hnt (1:10; DSHB 1G9, deposited by H.D. Lipshitz), Antp (1:100; DSHB 4C3, deposited by D. Brower), NimC1/P1 (1:300; I. Ando (Kurucz, Markus, et al., 2007)), Attila/L1 (1:100; I Ando (Kurucz, Vaczi, et al., 2007)), phospho-Histone H3Ser10 (1:500; Abcam ab14955). Lymph glands were next washed twice with PBT 0.2% and then incubated with fluorescent secondary antibodies for 2 hours in room temperature in the dark, followed by 10 min incubation with 1µg/mL Hoechst-33342 (Thermo Scientific™, Cat# 62249, Lot# PK1922313). The secondary antibodies used in this study were anti-mouse Alexa568 (1:1000, Molecular Probes, Cat# A10037) and anti-rat DyLight649 (1:1000, Jackson Immunoresearch, Cat#712-495153, Lot#93224). After staining, lymph glands were washed twice in PBT 0.2%, detached from the carcass and mounted on microscope slides with 15 µL ProLong™ Diamond Antifade Mountant (Invitrogen™, Cat#P36961). Typically, between 10 and 25 lymph glands were imaged and analyzed per genotype. Confocal imaging was performed on a Nikon Yokogawa CSU-W1 with a S Plan-FI LWD 20x air objective and a laser set of 405 nm, 488 nm and 561 nm. Lymph glands were imaged in a z-stack covering 25 µm with a z-step of 0.1 µm.

### Immunofluorescence of circulating hemocytes

For confocal imaging of circulating hemocytes, larvae were bled in ice-cold ESF921 media (Expression Systems, Cat# 500302) and the hemolymph from one larva was transferred to a corresponding well in a black 384-well glass-bottom plate (CellVis, Cat# P384-1.5H-N) The circulating hemocytes were fixed with 4% paraformaldehyde (Polysciences, Cat# 18814-20) and centrifuged for 5 min at 1000 rpm before incubation for 10 min at room temperature. Wells were then washed twice with PBT 0.2% and blocked with PBT 0.2% containing 5% bovine serum albumin (Roche, Cat# 10735094001). Primary antibody staining with anti-Wg (1:500, DSHB 4D4, deposited by S. M. Cohen) was done for 1 hour at room temperature, followed by 3 washes of PBT 0.2%. Wells were incubated with Alexa Fluor™ 647 Phalloidin (1:100, Molecular Probes, Cat#A22287) together with secondary antibody anti-mouse Alexa Fluor™ 568 (1:1000, Molecular Probes, Cat# A10037) and 1 µg/mL Hoechst-33342 (Thermo Scientific™, Cat# 62249, Lot# PK1922313) for 1 hour at room temperature, washed once in PBS and then mounted with 15 µL ProLong™ Diamond Antifade Mountant (Invitrogen™, Cat#P36961) per well. Imaging was done with Nikon Yokogawa CSU-W1 with S Plan-FI LWD 20x air objective where 5 images per larva was acquired.

### RT-qPCR

RT-qPCR was used to determine RNA levels of genes of interest, as well as to validate knockdown efficiency of RNAi fly lines. For qPCR on circulating hemocytes, the hemocytes were extracted by a protocol adapted from Tattikota and Perrimon (Tattikota & Perrimon, 2021). For each biological replicate, 15 larvae were collected in a 2 mL Eppendorf tube containing 500 µl glass beads and 200 µl PBS. The tube was vortex for 2 min at 2000 rpm. Only larvae where the heartbeat was detectable were included in the following. Each larva was dried and placed in a 10µl drop of ice-cold PBS before peeling the cuticle open allowing the hemocytes to enter the media by diffusion. The next larva was added to the same drop and the procedure repeated until all larvae were processed before the hemolymph was transferred to a new tube. RNA was extracted using TRIzol (Life Technologies, Cat#15596026, Lot#16200701) and the Zymo Research Direct-zol™ RNA MicroPrep kit (Cat# R2062, Lot# ZRC201718) and RNA concentration measured by NanoDrop. RNA was converted to cDNA using the iScript™ cDNA Synthesis Kit (BIO-RAD Cat#1708891) on the Applied Biosystems SimpliAmp PCR machine. RT-qPCR was performed with Fast SYBRGreen Master Mix (Applied Biosystems, Cat#4385614) and run on Applied Biosystems StepOnePlus Real Time PCR system with StepOne v2.3 software. Efficiency of primer sets was tested before running expression level analysis. qPCR primers were selected from http://www.flyrnai.org/flyprimerbank (Hu et al., 2013). All primers used are listed in Supplementary Table 2.

### Flow cytometry of live lymph gland cells

Flow cytometry was used to determine lymph gland size and differentiation through levels of mature GFP-positive hemocytes in the lymph gland. Around 15 lymph glands were dissected out and trypsinized with 100 µL of 1x trypsin with EDTA without phenol red (Life Technologies™, Cat#15400054) for 15 minutes at 25 °C. Each sample was pipetted up and down 40 times to ensure proper tissue dissociation. The reaction was quenched with 300 µL of PBS with 2% FBS, before the sample was filtrated in FACS tubes with 35 µm filter cap (Falcon®, Cat# 352235). 1 µL of 1,677 mg/mL Propidium Iodide (Sigma Aldrich, Cat# P4170) was added right before analysis to identify live and dead cells. Samples were run on LSRII UV laser flow cytometer and the analysis was performed with FACSDiva and FlowJo software.

### Drug treatment

SGC0946 (Selleckchem, Cat# S7079 and MedChemtronica Cat# HY-15650/CS-3531, Lot#58947), i-BET (Merck Millipore, Cat# 401010) and MI-463 (Selleckchem Cat# S7816) were dissolved in DMSO as a stock solutions and added to molten (∼40 °C) standard fly food mixed with blue food colorant (Panduro, Cat# 1018108-104) at final concentrations of 5 uM, 10 uM and 20 uM of the drug and 0.1% DMSO. DMSO concentration was kept equal across all experimental vials. The exception was for the preliminary drug experiments in Supplementary Figure 6, where DMSO concentration varied and DMSO only controls were provided for each drug concentration. 5 mL of food was added to each vial and allowed to solidify. Crosses were initially performed on normal fly food with yeast paste before they were flipped onto vials with drugs 24 hours after crossing to ensure sufficient egg laying and fertilization. Incubation times and temperature conditions were as in regular crosses described above. The presence of blue color in the larvae gut ensured exposure to the food mixed with drug.

### Image processing and quantification

Within each set of experiments, images were captured with identical settings below pixel value saturation and post-processed identically. Intensity enhancements for visualization purposes on confocal microscopy images was performed using Adobe Photoshop, ImageJ or NIS Elements, and all intensity alterations were done equally for all genotypes to ensure comparativeness within each experiment. Confocal images shown in figures are maximum intensity projections made using the NIS Elements software. For confocal imaging performed on the Nikon Yokogawa CSU-W1 microscope, lymph gland volume and cell type volume fractions were calculated using the NIS Elements software. Quantifications for circulating hemocytes were also performed with NIS Elements. Phospho-H3 positive cell quantifications were done using ImageJ and is reported per lymph gland area to account for size differences. Quantifications of NimC1/P1 positive cells were performed with NIS Elements software and counted in 1 Z-plane visualizing the surface plane only due to improper antibody penetration of lymph gland tissue.

### Experimental design and statistics

Samples were not masked or randomized during data collection or analysis. Statistical analysis was performed using GraphPad Prism 5. The data was assumed to be normally distributed. For multiple comparisons, one-way ANOVA with Bonferroni post-testing was used. For single comparisons, unpaired two-sided Student’s t-test was performed.

## Author contributions

Julie Aarmo Johannessen: Formal analysis, Investigation, Writing—original draft, Writing— review & editing, Visualization

Miriam Formica: Formal analysis, Investigation, Writing—review & editing

Miriam Aarsund: Investigation, Writing—review & editing,

Nora Rojahn Bråthen: Investigation, Writing—review & editing,

Amani Al Outa: Investigation, Writing—review & editing

Marc Therrien: Writing—review & editing, Supervision

Jorrit Enserink: Writing—review & editing, Supervision

Helene Knævelsrud: Conceptualization, Formal analysis, Investigation, Writing—original draft, Writing—review & editing, Supervision, Project administration, Funding acquisition

## Acknowledgements

The authors would like to thank I. Ando, and R. Paro for generously sharing reagents. Stocks obtained from the Bloomington Drosophila Stock Center (NIH P40OD018537) and Vienna Drosophila Resource Center (VDRC) were used in this study. Several antibodies were obtained from the Developmental Studies Hybridoma Bank created by the NICHD of the NIH and maintained at The University of Iowa. The core facilities for Advanced Light Microscopy and Flow Cytometry at Oslo University Hospital, Gaustad and Montebello nodes, are acknowledged for access, help and services. H.K. was supported by fellowship 2017062 from the South-Eastern Norway Regional Health Authority and grant 30078 from the Research Council of Norway. The research leading to these results has received funding from the European Union’s Horizon 2020 research and innovation programme under the Marie Skłodowska-Curie grant agreement No 801133. This work was partly supported by the Research Council of Norway through its Centres of Excellence funding scheme, project number 262652. J.M.E. is supported by the Norwegian Health Authority South-East, grant numbers 2017064, 2018012 and 2019096; the Norwegian Cancer Society, grant numbers 182524 and 208012; and the Research Council of Norway grant numbers 261936 and 294916. M. T. was supported by Foundation Grant FDN-388023 from the Canadian Institute of Health.

## Supplementary figures

**Supplementary figure 1:**
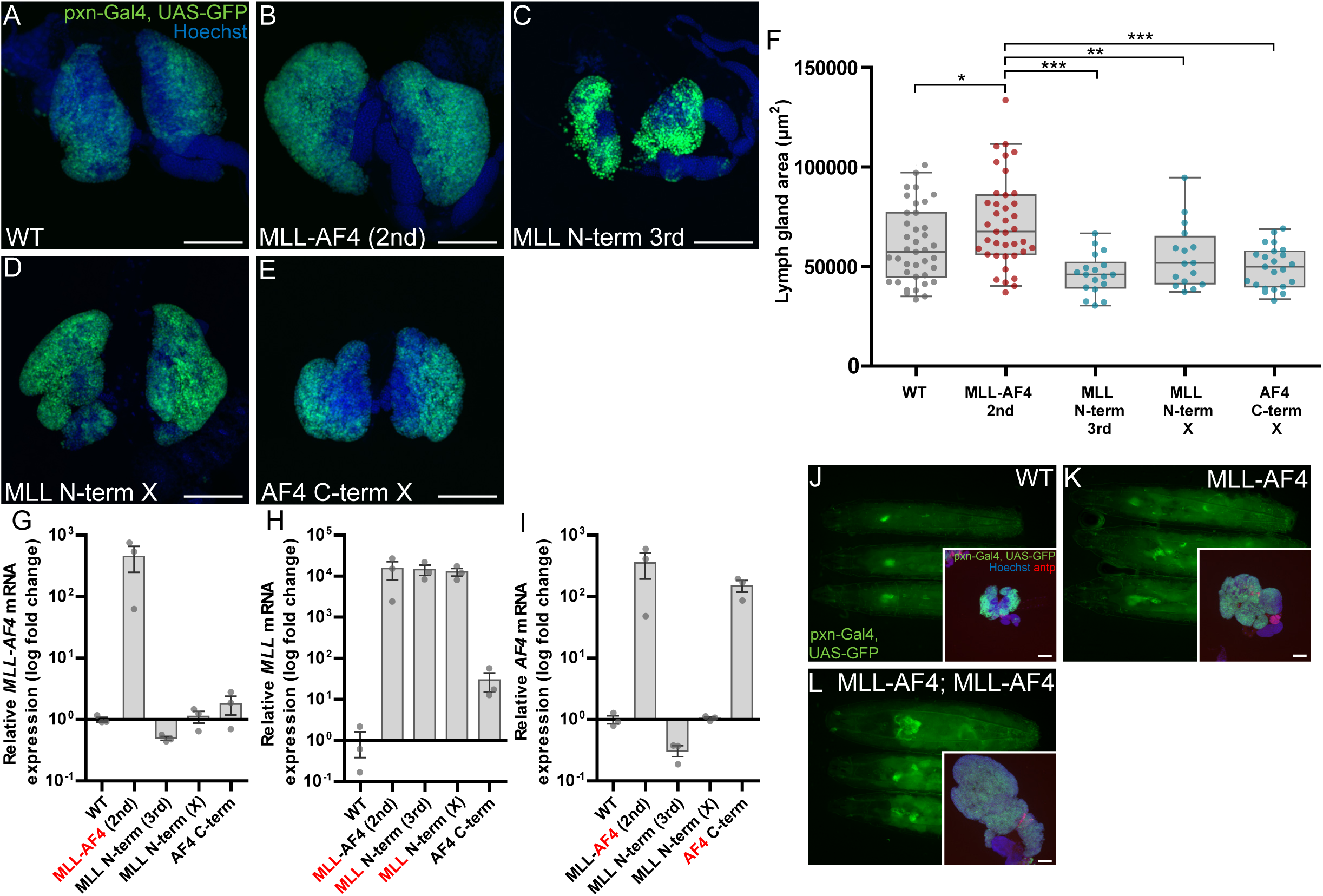
Expression of the N-terminal part of MLL or the C-terminal part of AF4 does not induce lymph gland hyperplasia. **A-E:** Immunofluorescence confocal images of lymph glands expressing human MLL-AF4, the N-terminal part or the C-terminal part of the MLL-AF4 transgene in the cortical zone. Cortical zone is marked by GFP expression. For MLL N-term, images are shown for transgenes on the 3^rd^ and the X chromosome. Images are maximum intensity projection of Z-stacks. Scale bars 100 µm. **F**: Quantification of lymph gland area (µm^2^) from immunofluorescence confocal images of lymph glands expressing either MLL-AF4 transgenes, the N-terminal MLL part or the C-terminal AF4 part of the fusion protein. Error bars of box plots show the 95^th^ percentile. One-way ANOVA with Bonferroni post-test was performed to assess significant differences. **G-I**: Expression levels of MLL-AF4 transgenes as measured by RT-qPCR. The part of the transgene that is detected by the qPCR primer set is highlighted in red. Experiments were conducted in three biological replicates and in technical duplicates, and values are normalized to relative expression in WT larvae. Error bars show standard error of the mean (SEM). **A-I:** 2^nd^, 3^rd^ and X refers to chromosome of transgene loci. **J-K:** Representative images of whole wandering third instar larvae expressing full-length human MLL, human MLL-AF4 on 2^nd^ chromosome or larvae expressing MLL-AF4 transgenes on both 2^nd^ and 3^rd^ chromosome driven by *pxn*-Gal4 imaged by widefield fluorescence microscopy. Inserts: Representative immunofluorescence confocal images of lymph glands. Scale bar 100 µm. **Genotypes: A, F, J**: *pxn-Gal4, UAS-GFP/+* (WT) **B, F, K**: *pxn-Gal4, UAS-GFP/UAS-MLL-AF4* **C, F:** *pxn-Gal4, UAS-GFP/+;UAS-MLL-N-term/+* **D, F***: UAS-MLL-N-term/+; pxn-Gal4, UAS-GFP/+* **E, F:** *UAS-AF4 C-term/+; pxn-Gal4, UAS-GFP/+* **G-I:** *hml-Gal4, UAS-GFP/+* (WT). *hml-Gal4, UAS-GFP/UAS-MLL-AF4* (UAS-MLL-AF4 (2^nd^)). *hml-Gal4, UAS-GFP/+;UAS-MLL-N-term/+* (UAS-MLL N-term (3^rd^)). *UAS-MLL-N-term/+; hml-Gal4, UAS-GFP/+* (UAS-MLL N-term (X)). *UAS-AF4 C-term/+; hml-Gal4, UAS-GFP/+* **L**: *pxn-Gal4, UAS-GFP/UAS-MLL-AF4; UAS-MLL-AF4/+*

**Supplementary figure 2:**
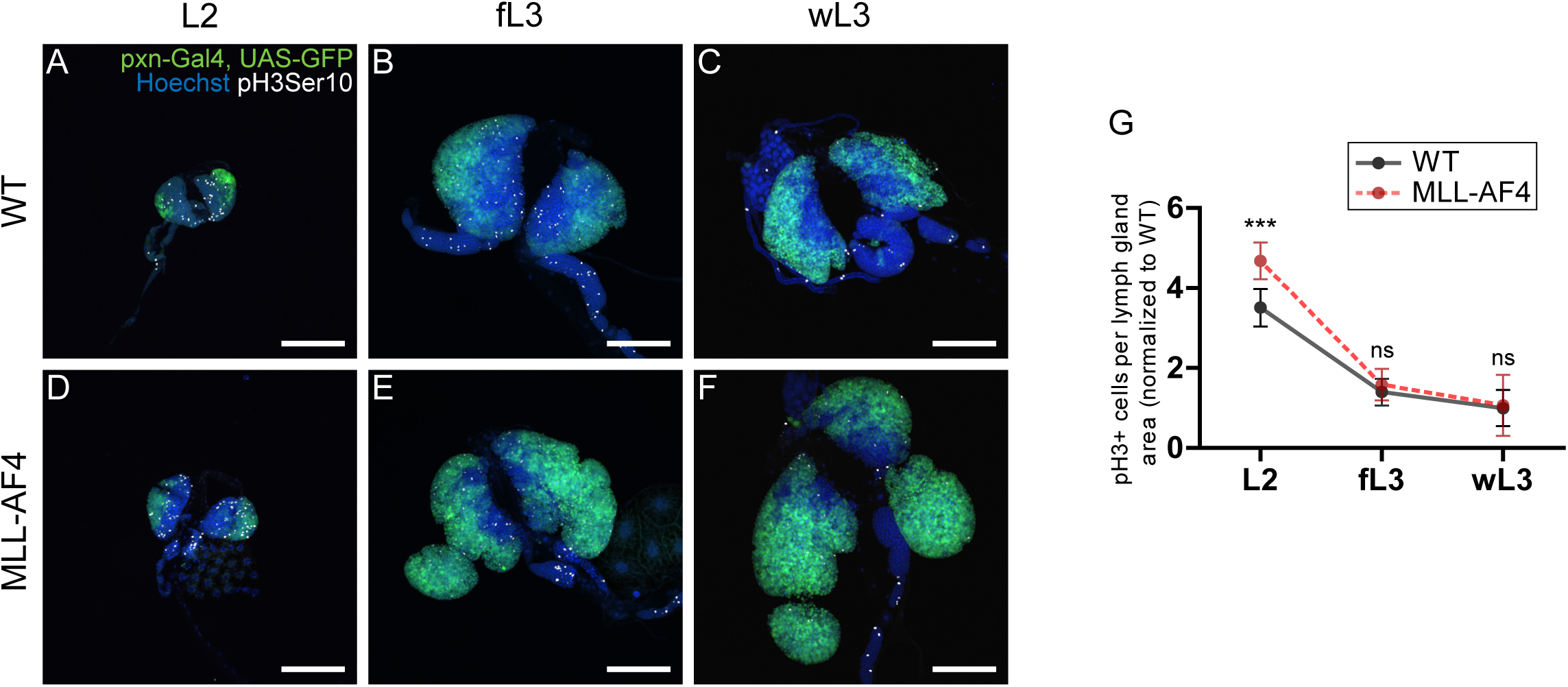
MLL-AF4 lymph glands have elevated levels of cell proliferation at an early larval stage. **A-F:** Immunofluorescence confocal images of lymph glands from larvae that are WT or expressing MLL-AF4 driven by *pxn*-Gal4, in 2^nd^ instar larva, feeding 3^rd^ instar larva and wandering 3^rd^ instar larva, respectively. Cortical zone is marked by GFP expression and phospho-histone 3 Ser10 (pH3Ser10) positive cells are shown in greyscale. Images are maximum intensity projection of Z-stacks. Scale bars 100 µm. **G:** Quantification of phospho-histone 3 Ser10 positive cells per lymph gland area for L2, fL3 and wL3 larval stages. All values are normalized to WT mean of wL3 larval stage. Error bars show 95^th^ percentile. Two-sided Student’s t-test was performed to assess significant differences between WT and MLL-AF4 for each larval stage. **Genotypes: A-C, G:** *pxn-Gal4, UAS-GFP/+* (WT), **D-F, G:** *pxn-Gal4, UAS-GFP/UAS-MLL-AF4*

**Supplementary figure 3:**
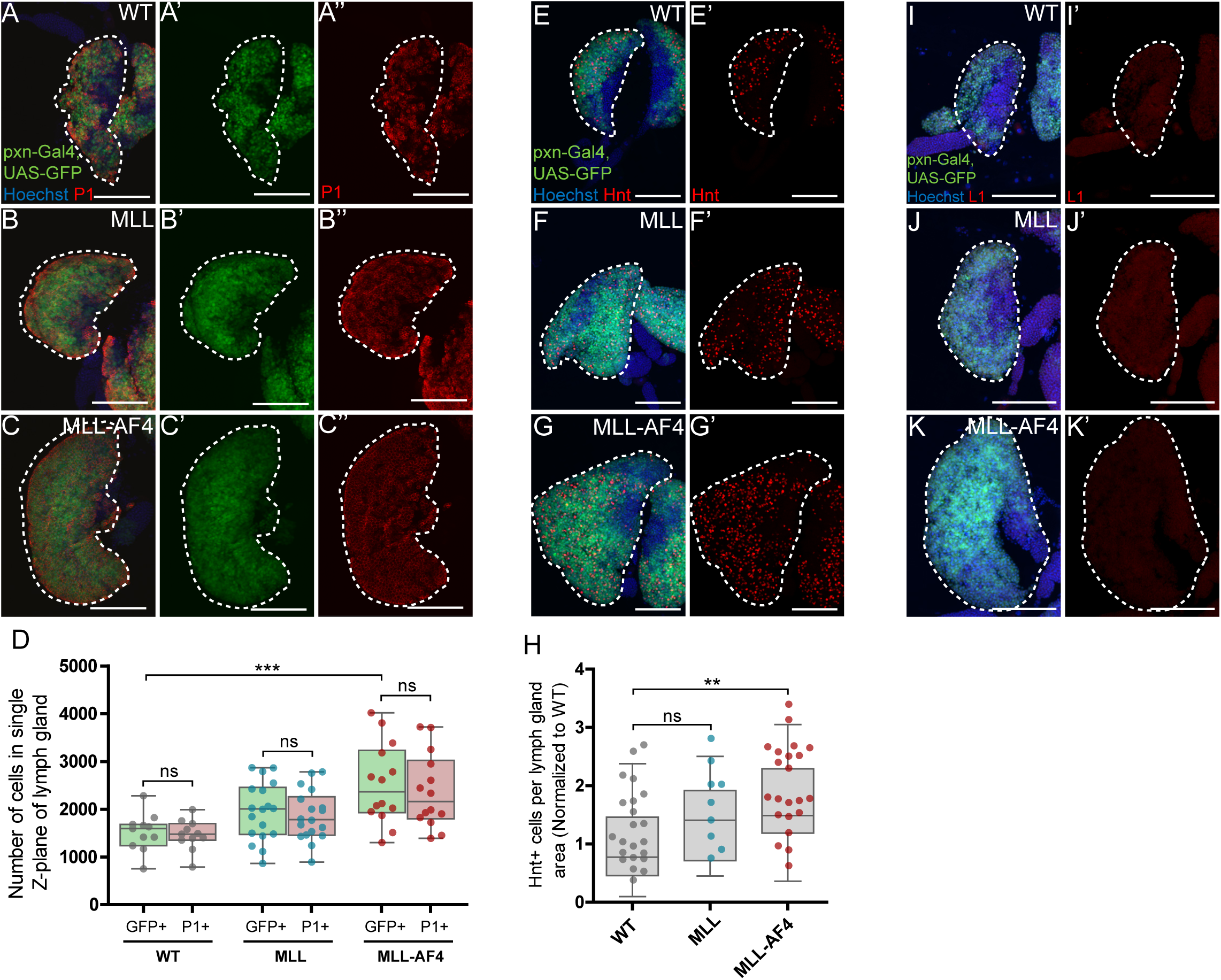
Characterization of the effects of MLL-AF4 on different hemocyte cell types. **A-C:** Immunofluorescence confocal images of lymph glands. The cortical zone is marked by GFP expression driven by *pxn*-Gal4 (A’-C’) and plasmatocytes are detected by staining against NimC1/P1 (shown in red, A’’-C’’). One primary lobe is outlined per genotype. **D:** Quantification of P1+ cells per GFP+ cell in lymph gland from immunofluorescence confocal images in A-C. **E-G:** Immunofluorescence confocal images of lymph glands. Cortical zone is marked by GFP expression driven by *pxn*-Gal4 and crystal cells are detected by staining against Hnt (shown in red, E’-G’). One primary lobe is outlined per genotype. Scale bars 100 µm. **H:** Quantification of Hnt+ cells per lymph gland from immunofluorescence confocal images in E-G. **I-K:** Immunofluorescence confocal images of lymph glands. Cortical zone is marked by GFP expression driven by *pxn*-Gal4 and lamellocytes are detected by staining against L1 (shown in red, I’-K’). One primary lobe is outlined per genotype. Scale bars 100 µm. **D, H:** Error bars of box plots show the 95^th^ percentile. One-way ANOVA with Bonferroni post-test was performed to assess significant differences. **Genotypes: A, D, E, H, I:** *pxn-Gal4, UAS-GFP/+* (WT) **B, D, F, H, J**: *pxn-Gal4, UAS-GFP/UAS-MLL* **C, F, G, H, K:** *pxn-Gal4, UAS-GFP/UAS-MLL-AF4*

**Supplementary figure 4:**
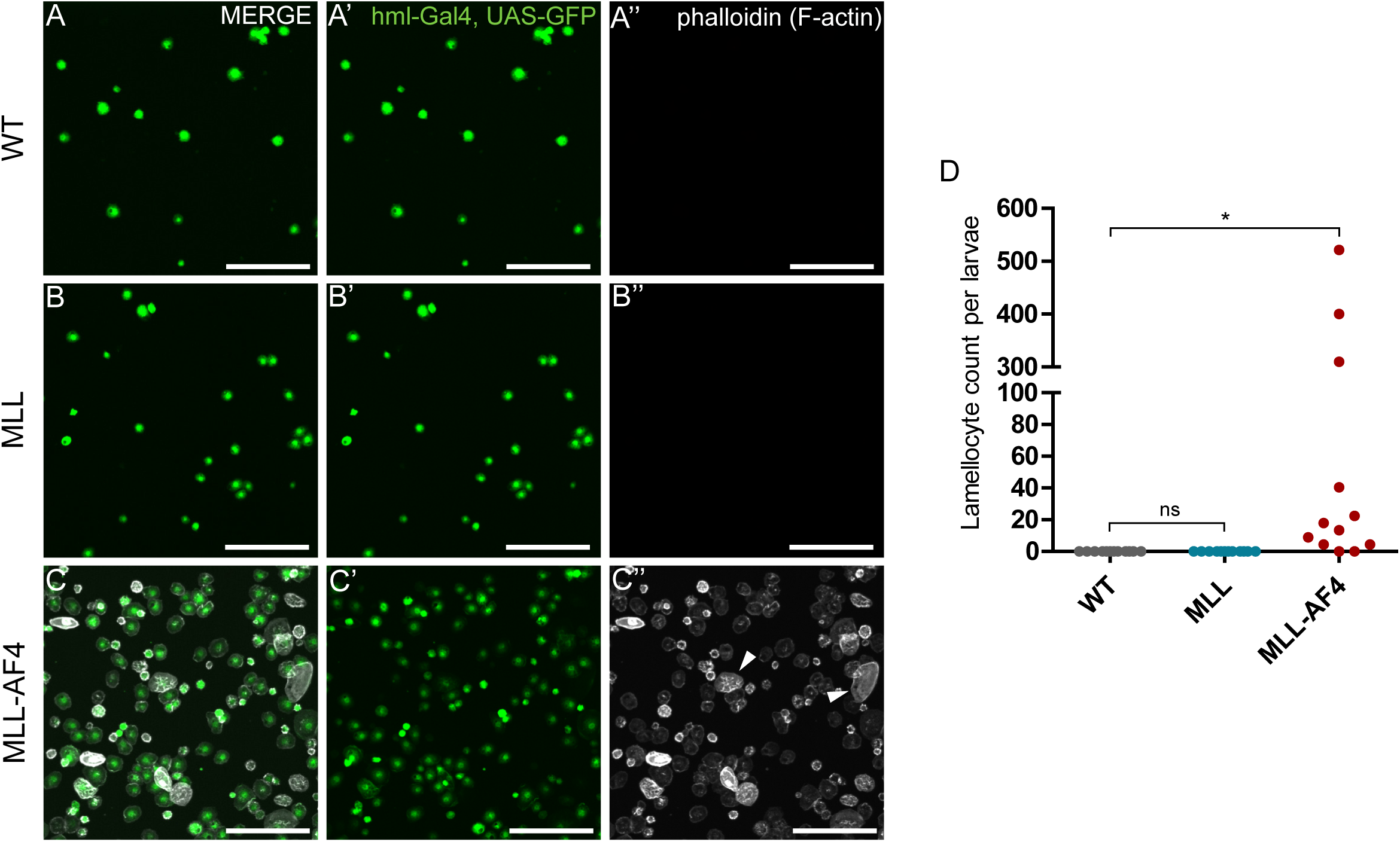
The increased number of circulating hemocytes consists of differentiated mature hemocytes and lamellocytes. **A-C:** Immunofluorescence confocal images of circulating hemocytes from larvae that are WT or expressing MLL-AF4 driven by *hml*-Gal4. Images are maximum intensity projection of Z-stacks. Scale bars 100 µm. A’-C’: GFP+ cells marked by the driver *hml-*Gal4, UAS-GFP. A’’-C’’: Phalloidin (F-actin) staining to visualize cytoskeleton and lamellocytes. Arrowheads indicate examples of differentiated lamellocytes. **D:** Quantification of lamellocytes in circulating hemocytes from A-C. Error bars of box plots show the 95^th^ percentile. One-way ANOVA with Bonferroni post-test was performed to assess significant differences. Note that the x-axis is split between 100 and 300 to enable visualization of all values. **Genotypes: A, D:** *pxn-Gal4, UAS-GFP/+* (WT) **B, D**: *pxn-Gal4, UAS-GFP/UAS-MLL* **C, D:** *pxn-Gal4, UAS-GFP/UAS-MLL-AF4*

**Supplementary figure 5:**
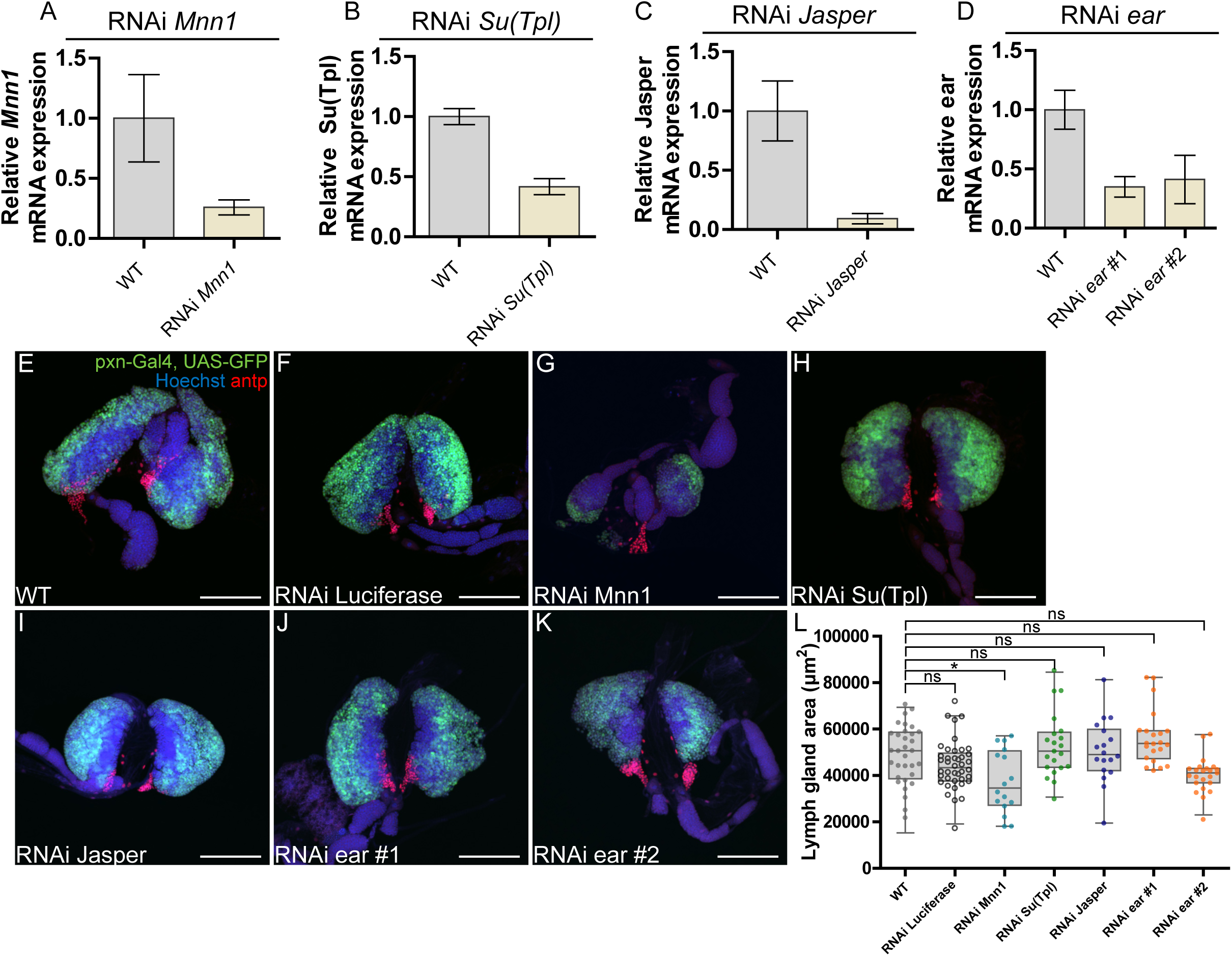
Expression of RNAi lines by themselves does not affect lymph gland size. **A-D:** Validation of knock-down efficiency through RT-qPCR measurements of relative expression of target genes. Error bars show the standard error of the mean (SEM). Complete overview over qPCR data for RNAi lines can be found in Supplementary Table 3. **E-K:** Representative immunofluorescence confocal images of lymph glands expressing RNAi targeting *Mnn1*, *Su(Tpl)*, *ear* and *Jasper* driven by *pxn-*Gal4. Cortical zone is marked by GFP expression and PSC is detected by immunostaining for antp (in red). Images are maximum intensity projection of Z-stacks. Scale bars 100 µm. **L:** Quantification of lymph gland area (µm^2^) from immunofluorescence confocal images of RNAi lines shown in E-K. Error bars of box plots show the 95^th^ percentile. One-way ANOVA with Bonferroni post-test was performed to assess significant differences. **Genotypes: A-D:** *hml-Gal4, UAS-GFP/+* (WT) **A:** *hml-Gal4, UAS-GFP/UAS-MLL-AF4; UAS-RNAi Mnn1 GL00018/+* **B:** *hml-Gal4, UAS-GFP/UAS-MLL-AF4; UAS-RNAi Su(Tpl) HMS00277/+* **C:** *hml-Gal4, UAS-GFP/UAS-RNAi Jasper HMC03961; UAS-MLL-AF4/+* **D**: *hml-Gal4, UAS-GFP/UAS-MLL-AF4; UAS-RNAi ear HMS00107/+* (#1) and *hml-Gal4, UAS-GFP/UAS-MLL-AF4; UAS-RNAi ear JF02905/+* (#2) **E, L:** *pxn-Gal4, UAS-GFP/+* (WT) **F, L:** *pxn-Gal4, UAS-GFP/+; UAS-RNAi Luciferase/+* **G, L**: *pxn-Gal4, UAS-GFP/UAS-MLL-AF4; UAS-RNAi Mnn1 GL00018/+* **H, L:** *pxn-Gal4, UAS-GFP/UAS-MLL-AF4; UAS-RNAi Su(Tpl) HMS00277/+* **I, L:** *pxn-Gal4, UAS-GFP/UAS-RNAi Jasper HMC03961; UAS-MLL-AF4/+* **J, L**: *pxn-Gal4, UAS-GFP/UAS-MLL-AF4; UAS-RNAi ear HMS00107/+* (#1) **K, L**: *pxn-Gal4, UAS-GFP/UAS-MLL-AF4; UAS-RNAi ear JF02905/+* (#2)

**Supplementary figure 6:**
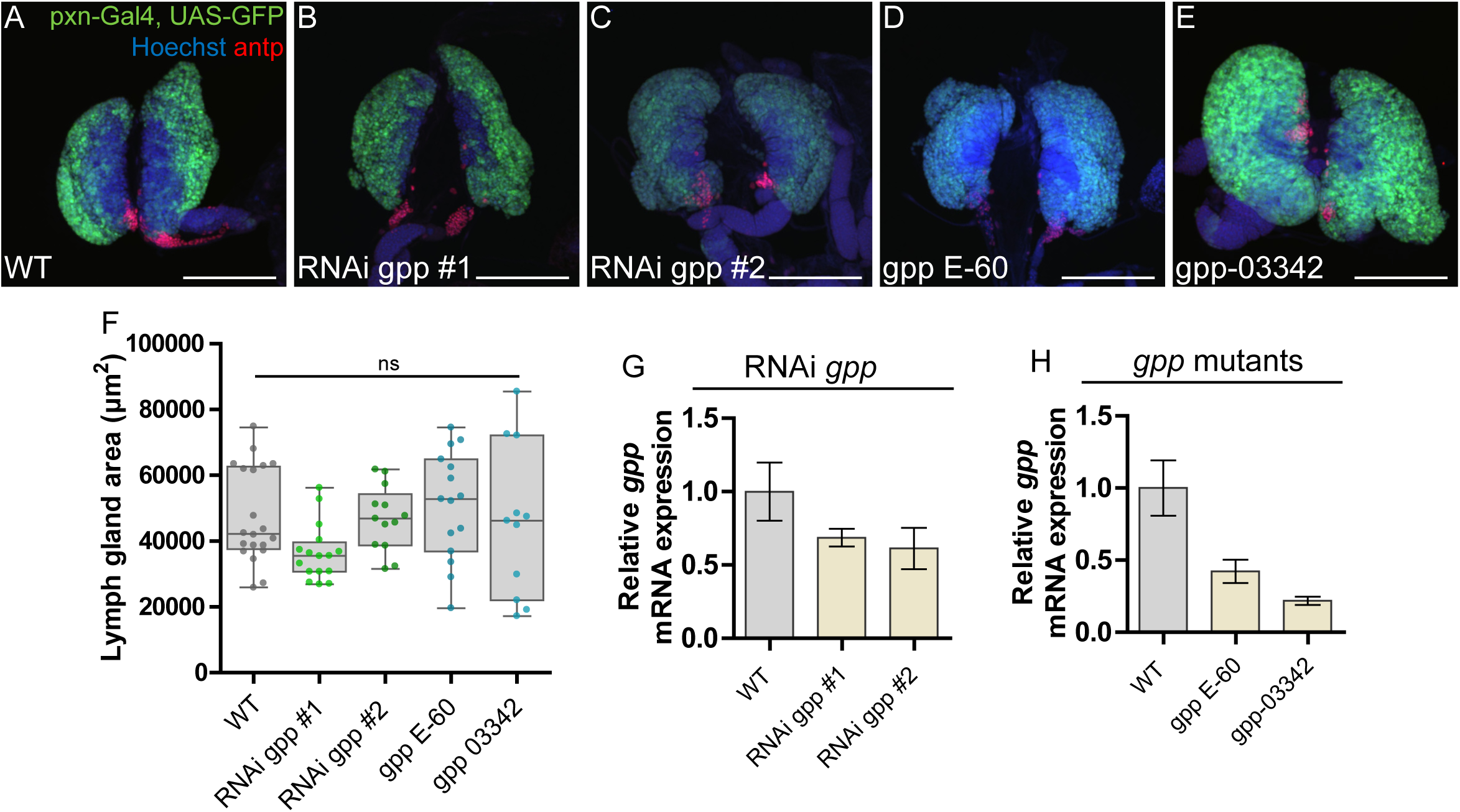
*gpp* RNAi and *gpp* mutant alleles do not affect the lymph gland phenotype. **A-E:** Representative immunofluorescence confocal images of lymph glands with reduced *gpp* levels through RNAi driven by *pxn*-Gal4 or introduction of *gpp* mutant alleles. Cortical zone is marked by GFP expression and PSC is detected by immunostaining for antp (in red). Images are maximum intensity projection of Z-stacks. Scale bars 100 µm. **F:** Quantification of lymph gland area (µm^2^) from immunofluorescence confocal images in A-E. Error bars of box plots show the 95^th^ percentile. One-way ANOVA with Bonferroni post-test was performed to assess significant differences. **G, H:** Validation of knock-down through RT-qPCR measurements of relative *gpp* expression levels in circulating hemocytes for RNAi lines and *gpp* alleles, respectively. Error bars show the standard error of the mean (SEM). Complete overview over qPCR data for RNAi lines can be found in Supplementary Table 3. **Genotypes: A, F:** *pxn-Gal4, UAS-GFP/+* (WT) **B, F:** *pxn-Gal4, UAS-GFP/+; UAS-RNAi gpp HMS00160/+* (#1) **C, F:** *pxn-Gal4, UAS-GFP/+; UAS-RNAi gpp JF1283/+* (#2) **D, F:** *pxn-Gal4, UAS-GFP/+; UAS-gpp E-60/+* **E, F:** *pxn-Gal4, UAS-GFP/+; UAS-gpp 03342/+* **G, H:** *hml-Gal4, UAS-GFP/+* (WT) **G:** *hml-Gal4, UAS-GFP/+; UAS-RNAi gpp HMS00160/+* (#1) and *hml-Gal4, UAS-GFP/+; UAS-RNAi gpp JF1283/+* (#2) **H:** *hml-Gal4, UAS-GFP/+; UAS-gpp E-60/+* and *hml-Gal4, UAS-GFP/+; UAS-gpp 03342/+*

**Supplementary figure 7:**
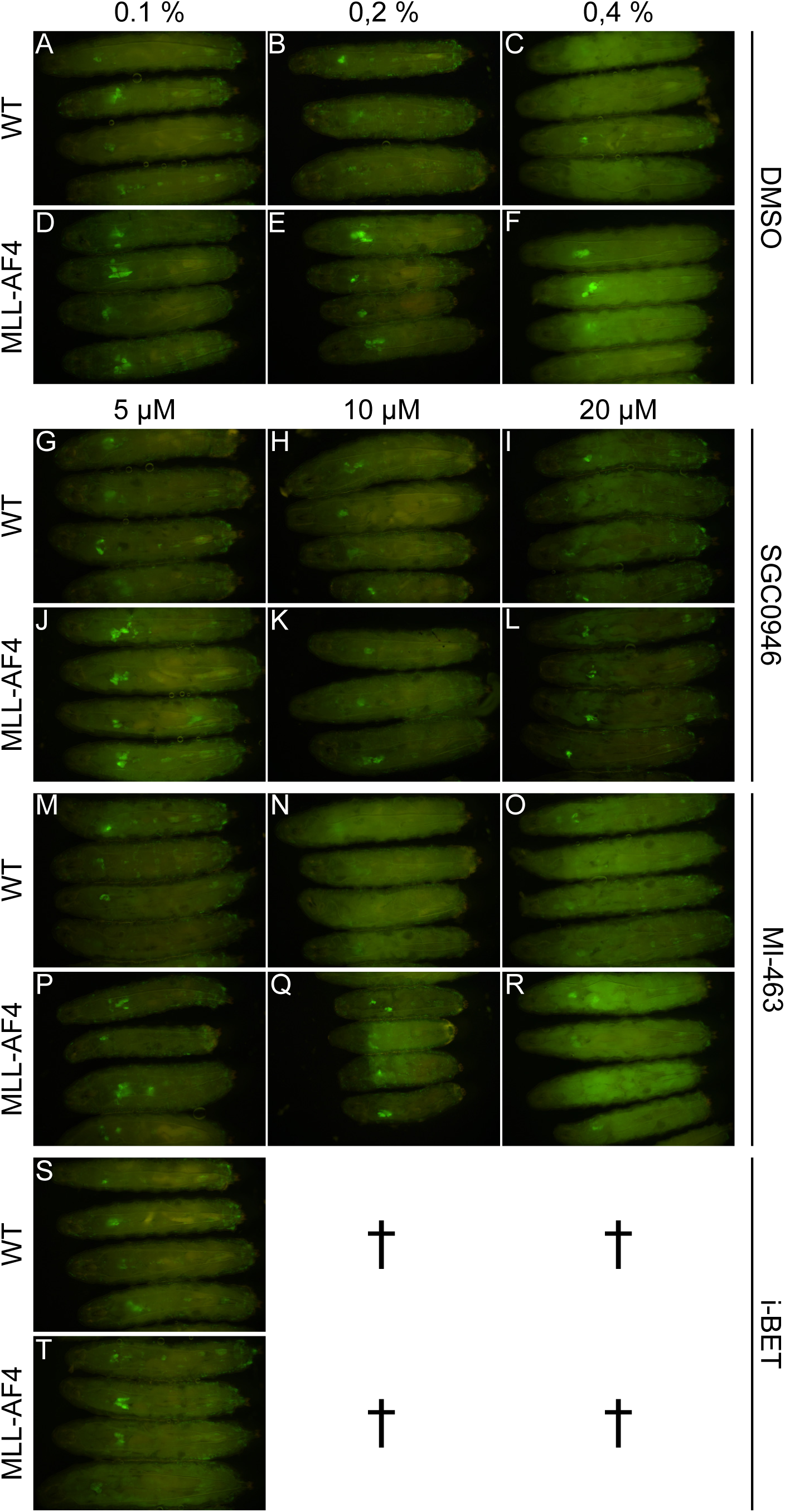
Drug treatment of larvae expressing MLL-AF4. **A:** Representative images of whole wandering third instar larvae treated with SGC0946, MI-643 and i-BET respectively in doses 5, 10 and 20 µM imaged by widefield fluorescence microscopy. Larvae are either WT or expressing human MLL-AF4 driven by *pxn*-GFP. The hematopoietic system is marked by GFP expression (*pxn*-Gal4). †: No live larvae after drug treatment. **Genotypes: A-C, G-I, M-O, S:** *pxn-Gal4, UAS-GFP/+* (WT) **D-F, J-L, P-R, T:** *pxn-Gal4, UAS-GFP/UAS-MLL-AF4*

**Supplementary table 1:** Fly lines

| Genotype | Reference in manuscript | Provider | Stock number/reference |
| --- | --- | --- | --- |
| w, pxn-Gal4, UAS-GFP |  | F. Lemieux | (Stramer et al., 2005) |
| wiso | WT | M. Therrien |  |
| w; UAS-MLL-AF4 ML5 (on 2<sup>nd</sup>) | MLL-AF4 (on 2<sup>nd</sup>) | R. Paro | (Muyrers-Chen et al., 2004) |
| w; UAS-MLL-AF4 ML6 (on 3<sup>rd</sup>) | MLL-AF4 (on 3<sup>rd</sup>) | R. Paro |  |
| w; UAS-MLL FL on 2nd |  | R. Paro |  |
| w; UAS-MLL FL on 3rd |  | R. Paro |  |
| w; UAS-MLL N-term (on 3<sup>rd</sup>) |  | R. Paro |  |
| w; UAS-MLL N-term (on X) |  | R. Paro |  |
| w; UAS-AF4 C-term on X |  | R. Paro |  |
| UAS-RNAi ear TRiP.HMS00107}attP2 | RNAi ear #2 | Bloomington | 62249 |
| UAS-RNAi ear TRiP.JF02905}attP2 | RNAi ear #1 | Bloomington | 56376 |
| UAS-RNAi Su(Tpl) TRiP.GL00168}attP2 |  | Bloomington | 35270 |
| UAS-RNAi Su(Tpl) TRiP.HMS00277}attP2 | RNAi Su(Tpl) | Bloomington | 33399 |
| UAS-RNAi Mnn1 TRiP.HMC03436}attP40 |  | Bloomington | 51862 |
| UAS-RNAi Mnn1 TRiP.JF01775}attP2 |  | Bloomington | 31220 |
| UAS-RNAi Mnn1 TRiP.GL00018}attP2 | RNAi Mnn1 | Bloomington | 35150 |
| w[1118]; UAS-RNAi Mnn1 GD8117 |  | VDRC | 17701 |
| UAS-RNAi Mnn1 KK101050 |  | VDRC | 110376 |
| UAS-RNAi-CG7946 TRiP.HMC03961}attP40 | RNAi Jasper | Bloomington | 55274 |
| y[1] v[1]; UAS-gpp-RNAi (TRiP.JF01284) (on 3rd) |  | Bloomington | 31327 |
| y[1] v[1]; UAS-gpp-RNAi (TRiP.JF01283) (on 3rd) | RNAi gpp #2 | Bloomington | 31481 |
| UAS-RNAi gpp VDRC 47199 |  | VDRC | 47199 |
| y[1] sc[*] v[1]; UAS-RNAi-gpp (TRiP.HMS00160) (on 3rd) | RNAi gpp #1 | Bloomington | 34842 |
| y[1] sc[*] v[1]; UAS-gpp-RNAi (TRiP.GL01325) (on 3rd) |  | Bloomington | 41893 |
| y[1] sc[*] v[1]; UAS-gpp-RNAi (TRiP.HMS02612) (on 2nd) |  | Bloomington | 42919 |
| UAS-gpp-RNAi KK107875 |  | VDRC | 110264 |
| gpp E-60/TM6b |  | M. Therrien | (Gavory et al., 2021) |
| gpp[03342]/TM6b |  | Bloomington | 11585 |
| y[1] v[1]; UAS-luciferase-RNAi TRiP.JF01355 |  | Bloomington | 31603 |
| w[1118]; P{w[+mC]=UAS-lacZ.NZ}J312 |  | Bloomington | 3956 |
| UAS-RNAi-lilli TRiP.JF02087}attP2 |  | Bloomington | 26314 |
| UAS-RNAi-lilli TRiP.HMS01066}attP2 |  | Bloomington | 34592 |

**Supplementary table 2:**
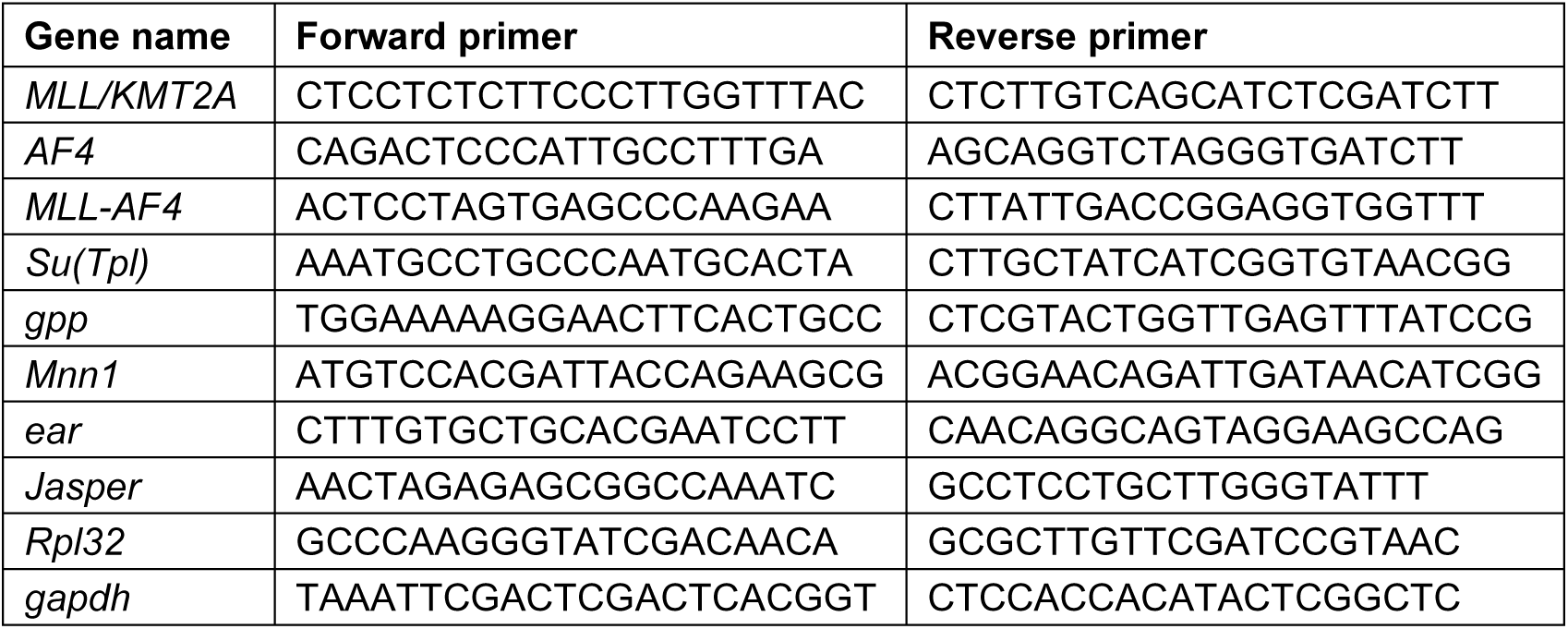
Primer sets used in RT-qPCR

**Supplementary table 3:** RT-qPCR results for all RNAi lines tested for knock-down efficiency in circulating hemocytes (hml-Gal4; UAS-GFP)

| Target | RNAi (or mutant) line | Mean relative expression of target gene (normalized to WT) | Standard deviation |
| --- | --- | --- | --- |
| Mnn1 | UAS-RNAi Mnn1 TRiP.GL00018}attP2 | 0,2591 | 0,1082 |
| Mnn1 | UAS-RNAi Mnn1 TRiP.JF01775}attP2 | 0,9281 | 0,5574 |
| Mnn1 | UAS-RNAi Mnn1 TRiP.HMC03436}attP40 | 0,0829 | 0,0525 |
| Mnn1 | w[1118]; UAS-RNAi Mnn1 GD8117 | 0,7720 | 0,0396 |
| Mnn1 | UAS-RNAi Mnn1 KK101050 | 0,4792 | 0,1686 |
| gpp | w[1118]; UAS-gpp-RNAi/CyO VDRC 47199 | 1,0582 | 0,2757 |
| gpp | y[1] v[1]; UAS-gpp-RNAi (TRiP.JF01284) (on 3rd) | 0,3778 | 0,1071 |
| gpp | y[1] v[1]; UAS-gpp-RNAi (TRiP.JF01283) (on 3rd) | 0,6124 | 0,2450 |
| gpp | y[1] sc[*] v[1]; UAS-RNAi-gpp (TRiP.HMS00160) (on 3rd) | 0,6864 | 0,1030 |
| gpp | y[1] sc[*] v[1]; UAS-gpp-RNAi (TRiP.GL01325) (on 3rd) | 0,8638 | 0,2574 |
| gpp | y[1] sc[*] v[1]; UAS-gpp-RNAi (TRiP.HMS02612) (on 2nd) | 0,6263 | 0,1941 |
| gpp | UAS-gpp-RNAi VDRC KK110264 | 0,5784 | 0,1126 |
| gpp | gpp E-60/TM6b | 0,4222 | 0,1385 |
| gpp | gpp[03342]/TM6b, Tb | 0,2185 | 0,0483 |
| Su(Tpl) | UAS-RNAi Su(Tpl) TRiP.GL00168}attP2/TM3, Sb[1] | 1,1418 | 0,2144 |
| Su(Tpl) | UAS-RNAi Su(Tpl) TRiP.HMS00277}attP2 | 0,4173 | 0,1145 |
| ear | UAS-RNAi ear TRiP.JF02905}attP2 | 0,7164 | 0,6151 |
| ear | UAS-RNAi ear TRiP.HMS00107}attP2 | 0,6065 | 0,2594 |
| Jasper | UAS-RNAi-CG7946 TRiP.HMC03961}attP40 | 0,1049 | 0,0494 |
| lilli | UAS-RNAi-lilli TRiP.JF02087}attP2 | 1,6282 | 0,3367 |
| lilli | UAS-RNAi-lilli TRiP.HMS01066}attP2 | 0,9807 | 0,3801 |

## Notes

### Competing Interest Statement

The authors have declared no competing interest.

